# Hypoxia-inducible factor 2-alpha promotes fitness of cytokine-activated human natural killer cells

**DOI:** 10.64898/2026.11.05.764649

**Authors:** H. Alex Feldman, Andrew VonHandorf, Kate E. Alianiello, Krista L. English, Seth D. Reighard, Hermine I. Brunner, Leah C. Kottyan, Matthew T. Weirauch, Kenneth M. Kaufman, Stephen N. Waggoner

## Abstract

Natural Killer (NK) cells are innate effector cells with potent antiviral, antitumor, and immunoregulatory functions that are relevant for the treatment of human diseases. There is a pressing need to better understand the mechanisms governing cellular fitness and functional persistence of NK cells to generate NK cell-based therapeutics of sufficient durability to achieve meaningful clinical benefit. Given the central role of interleukin-2 (IL-2) and IL-15 in the survival and proliferation of NK cells, we sought to identify transcription factors induced by these cytokines that are essential for maintaining the fitness of human NK cells. Culture in IL-2 or IL-15 increased protein expression of hypoxia inducible factor 2 alpha (HIF-2α, encoded by *EPAS1*) but not HIF-1α in NK cells from healthy human blood donors. Unexpectedly, induction of HIF-2α was independent of limiting oxygen. CRISPR targeting of *EPAS1* or pharmacological inhibition of HIF-2α protein expression markedly compromised the survival of cytokine-activated human NK cells. In contrast, inhibition of canonical HIF-2α transcription factor activity via blocking HIF-2α /HIF-1β heterodimer formation had no apparent effects on NK cell viability, function, or metabolic fitness. These results suggest that cytokines, rather than hypoxia, activate an unconventional HIF-1β independent, HIF-2α pathway that helps to maintain NK cells during cellular activation, revealing the potential promise of manipulating HIF-2α to improve the efficacy of adoptive NK cell therapies.

**Key points:**

- Cytokine activation of human NK cells induces hypoxia-independent HIF-2α protein expression
- Conventional HIF-2α/HIF-1β heterodimer driven gene regulation is dispensable in NK cells
- An unconventional activity of HIF-2α promotes survival of cytokine-activated NK cells.

## INTRODUCTION

Natural Killer (NK) cells are innate lymphocytes that play key roles in tumor surveillance, antiviral immunity, and immunoregulation (1-3). These functions are achieved through granule-mediated cytolytic killing of target cells and production of antiviral cytokines, including IFN-*γ* (4). Indeed, a variety of NK cell-based therapeutics for cancer and infection are progressing through clinical testing (5, 6). Activated and expanded NK cells derived from either patients (autologous) or unrelated sources (allogeneic), including products genetically engineered to express a chimeric antigen receptor (CAR) or other enhancements, have demonstrated safe and effective antitumor functions in patients (7, 8).

The initial efficacies of CAR-T and CAR-NK cells in killing of cancer cells are comparable, yet a widely perceived limitation of NK cells is the putatively inferior ability of these cells to durably persist following infusion (9, 10). While head-to-head clinical comparisons are lacking, the proposed ‘persistence deficit’ of NK cells is often ascribed to a reduced proliferative capacity and weaker functional resilience of NK cells within tumors when compared to T cells (11, 12). However, an alternative explanation is that our current knowledge and translational capacity to optimize the persistence of NK cells is limited in comparison to our understanding of T cell biology.

We sought to identify mediators that support the fitness and persistence of activated human NK cells. We focused on the cytokines interleukin-2 (IL-2) and IL-15, which are established regulators of growth, proliferation, and survival in NK cells (13). These and related cytokines are frequently used to expand NK cells *in vitro* and bolster cellular persistence once infused into patients (14, 15). We hypothesized that these cytokines induce mediators, some known and others yet to be identified, that enhance the fitness and durable function of activated NK cells.

In this study, CRISPR-Cas9 (clustered regularly interspaced short palindromic repeats) perturbation experiments in human NK cells identify hypoxia inducible factor 2 alpha (HIF-2α encoded by *EPAS1*) as a candidate IL-2/IL-15-responsive driver of NK cell fitness. Conventionally, hypoxic conditions stabilize the HIF-1α and HIF-2α proteins by preventing the rapid hydroxylation, ubiquitination, and degradation of these proteins when oxygen is plentiful (16, 17). Upon stabilization, either HIF-1α or HIF-2α can heterodimerize with HIF-1β and drive transcription of genes involved in cellular adaptation to hypoxia via binding to hypoxia responsive elements within the genome (18). There are overlapping and distinct genes regulated by HIF-1α and HIF-2α (19-21). Our current understanding of the role of HIF-2α in NK cell biology is limited. In tumor cells, HIF-2α modulates susceptibility to NK cell killing (22), but corresponding functions for HIF-2α in NK cells are as yet undescribed. In contrast, up-regulation of HIF-1α is a hallmark of the NK cell response to hypoxia. HIF-1α has been described to play both negative and positive roles in NK-cell survival, metabolic activity, cytolytic capacity, and inflammatory functions (23-32). These observations likely reflect the need to manage HIF-driven transcriptional responses in NK cells in the context of hypoxic tumor microenvironments.

Herein, we provide a demonstration of unconventional stabilization of HIF-2α, but not HIF-1α, in human NK cells following treatment with inflammatory cytokines. We further demonstrate a necessity for HIF-2α in cytokine-activated NK cell fitness that is independent of classical HIF-2α/HIF-1β-driven transcriptional activity.

## METHODS

### Isolation of NK cells from healthy blood donors

Leukocyte reduction filters from healthy adult blood donors were sourced from the Hoxworth Blood Center with an exemption designation from the Cincinnati Children’s Hospital Medical Center Institutional Review Board. To isolate leukocytes, filters were repeatedly washed with phosphate-buffered saline (PBS) before centrifugation separation using PBMC (peripheral blood mononuclear cell) Spin gradient centrifugation media in Plurimate II 50 mL tubes (PluriSelect, Leipzig, Germany). Following additional wash steps, residual red blood cells were lysed with Ammonium-Chloride-Potassium (ACK) buffer at 37°C for 5 minutes. Subsequent cell pellets were resuspended in MACS buffer (PBS, 0.5% FBS, 2 mM EDTA) for NK cell enrichment using magnetic bead-based negative selection kits (Miltenyi Biotec, Gaithersburg, MD). Enriched NK cells were resuspended in RPMI-1640 supplemented with 10% fetal bovine serum (FBS), Penicillin/Streptomycin, and L-Glutamine at 2 to 3 x 106 cells/mL (denoted as complete RPMI). Cells were rested overnight and then stimulated with 400 U/mL of recombinant human (rh) IL-2 (Peprotech, Cranbury, NJ) or 10 ng/mL rhIL-15 (Thermo Fisher, Waltham, MA), or 10 ng/mL of rhIL-12 and 100 ng/mL of rhIL-18 (Peprotech).

### Small molecule inhibitors

All small molecules were purchased from MedChemExpress (Monmouth Junction, NJ) unless otherwise noted. After resting overnight following enrichment, NK cells were treated with either HIF-2α heterodimerization inhibitor PT2385 (10 μM), HIF-2α Translation Inhibitor (C76, 10μM, Sigma-Aldrich, Burlington, MA), or the HIF-1α inhibitor (CAY10585, 20 μM, Cayman Chemical, Ann Arbor, MI) or HIF-1α translation inhibitor (KC7F2, 20 μM) dissolved in dimethyl sulfoxide (DMSO). In all experiments, cells treated with inhibitors were compared to cells treated with similar concentrations of DMSO alone.

### Culture of K562 Cells

K562 cells were acquired from ATCC (Manassas, VA). K562 cells were thawed from -80°C stocks and cultured in complete RPMI-1640 at a density of 1×10^6^ cells/mL. Cells were cultured in a 37°C incubator with 5% CO_2_. Media was refreshed every 3 days, and dead cells were eliminated via periodic spins with Ficoll (Cytiva, Marlborough, MA).

### Expansion of NK cells

NK cells were expanded using the CTS NK Expander system (Gibco, Waltham, MA) with slight modifications. Briefly, 2×10^6^ NK cells were resuspended in 2.5 mL of complete CTS NK Expansion media supplemented with 500 U/mL of human IL-2. After 5 days of expansion, cells were counted and diluted to 7.5 x 10^5^ cells/mL in CTS NK Expansion Media and given fresh IL-2. Cell density was assessed every 2 days afterwards. The above process was repeated for a maximum of 14 days of proliferation after initial seeding. After 14 days, remaining cells were washed once with sterile PBS and plated in complete RPMI-1640 at 2-3 ×10^6^ cells/mL and supplemented with 400 U/mL of IL-2. Expanded NK cells were used in Seahorse assays as outlined below.

### Quantitative (Real-Time) Polymerase Chain Reaction (qPCR)

RNA from enriched NK cells was isolated using the RNeasy Mini Kit (Qiagen, Germantown, MD) according to the manufacturer’s instructions. In most cases, the maximum cell number recommended by the manufacturer (1×10^6^ cells) was used. If not immediately making cDNA, RNA was stored at - 80°C. To make cDNA, 50 ng of RNA was combined with 4 μL iScript Reverse Transcriptase Supermix (BioRad, Hercules, CA) and nuclease-free water up to a final volume of 20 μL. cDNA was generated using the manufacturer’s recommended thermocycler program. For qPCR experiments, 1 part cDNA was diluted in 4 parts sterile H_2_O after verifying amplification and efficiency of reaction. For each well of a 96-well 0.1 mL Fast plate (Thermo Fisher), 2 μL diluted cDNA was combined with 10 μL TaqMan Fast qPCR Master Mix (Thermo Fisher) and 7 μL of sterile water before corresponding TaqMan probes were added. qPCR reactions were run on a QuantStudio 5 (Applied Biosystems, Waltham, MA) using the standard “Fast” protocol. All samples were run in triplicate. The following TaqMan probes were used: Hs01026142_m1 (*EPAS1*) and Hs02786624_g1 (*GAPDH*)

### Flow cytometry and Intranuclear Staining

Enriched NK cells were plated in a 96-well U bottom plate (Corning, Corning, NY) and centrifuged at 400 x *g* for 5 minutes. Cells were then washed with 200 μL of Hank’s Buffered Salt Solution without calcium or magnesium, supplemented with 1% FBS and 2mM EDTA to prevent cell clumping (FACS buffer). After another centrifugation step, cells were stained for 5 minutes at room temperature with Zombie NIR Live/Dead (Biolegend, San Diego, CA) diluted 1:1,000 in PBS. Cells were centrifuged and washed twice in 200 μL FACS buffer. Cells were stained with a combination of the following antibodies in FACS buffer supplemented with 10% human serum: CD3 (Clone: SK7; BD Biosciences, Milpitas, CA), CD14 (Clone: MφP9; BD Biosciences), CD19 (Clone: HIB19; BD Biosciences), HLA-DR (Clone: QA19A44; Biolegend), CD56 (Clone HCD56; Biolegend), or CD16 (Clone: 3G8; Biolegend). After 25 minutes at room temperature, cells were centrifuged and washed with FACS buffer before adding 100 μL of Cytofix (BD Biosciences) and incubating for 4 minutes at 4°C. Finally, cells were centrifuged and washed twice with FACS buffer before resuspending in 200 μL of FACS buffer. For intranuclear staining, 100 μL FoxP3 Transcription Factor (TF) Fix/Perm buffer (BD Biosciences) was added to cells, incubated for 45 minutes at 4°C, and washed with TF Fix/Perm according to the manufacturer instructions. Cells were stained with antibodies for HIF-1α (Clone: 546-16; Biolegend) and HIF-2α (Clone: ep190b; Novus Biologics, Centennial, CO) overnight (~16 hours) at 4°C. The next day, cells were centrifuged at 400x *g* for 6 minutes, washed once with TF Fix/Perm wash, once with FACS buffer, and resuspended in 200 μL FACS buffer for analysis. For cell death experiments we employed the Annexin V Apoptosis Detection Kit with PI (Miltenyi). Cells were stained with antibodies as above, washed once with FACS buffer, and resuspended in 100 μL of Annexin V binding buffer. Subsequently, 10 μL of Annexin V was added to each well, mixed, and incubated for 15 minutes at room temperature protected from light. Afterwards cells were washed twice with binding buffer before final resuspension in 500 μL of binding buffer. 5 μL of propidium iodide solution was added immediately preceding flow cytometry analysis. All cells were counted using a hemocytometer and the Trypan blue exclusion method. All flow cytometry experiments were run on an LSR Fortessa or FACSymphony A5 SE cytometer (BD Biosciences). All analyses were completed in FlowJo (FlowJo, Ashland, OR).

### NK cell killing assay

Cells were collected by vigorous pipetting and centrifuged. Subsequently, NK cells were counted and resuspended at 2 ×10^6^ cells/mL in complete RPMI-1640 supplemented with 50 U/mL of human IL-2. 200 μL of NK cells were added to a 96-well U bottom plate before serially diluting the NK cells 5 more times in 100 μL of media. K562 targets were resuspended in complete RPMI-1640 at 1×10^6^ cells/mL before adding Calcein AM dye (Thermo Fisher) to a final concentration of 3.33 μg/mL (1:300 dilution of Calcein AM stock). Cells were then placed in a 37°C incubator for 1 hour with occasional vortexing. Calcein labeling was confirmed visually by examining the change in color of the K562 pellet. After washing twice, K562 cells were diluted to 1×10^5^ cells/ mL and 100 μL of cells were added to all NK wells. Maximum lysis was defined as K562 cells plated with 2% Triton-X100 in RPMI-1640. Spontaneous release of Calcein was defined as the fluorescence of the supernatant from wells containing K562 cells but not effectors or detergents. After 4 hours, 100μL of supernatant was collected into a new 96-well plate and fluorescence was measured on a Glo-Max instrument (Promega, Madison, WI) with excitation filter 485 nm, emission filter 530 nm. % lysis was calculated as: [(test release-spontaneous release) / (maximum release - spontaneous release)] x 100. Killing assays were run in triplicate. Spontaneous or maximum release was averaged over at least 6 wells.

### NK cell cytokine secretion

Following isolation, cells were rested overnight before stimulation with 10 ng/mL of human IL-12 and 100 ng/mL of human IL-18 (Peprotech) for 16 hours. GolgiStop solution was made by adding 4 μL of GolgiStop (monensin; BD Biosciences) to 2 μL of sterile water. After 16 hours, 1 μL of GolgiStop solution and 1μL of GolgiPlug (brefeldin A; BD Biosciences) were added per 1 mL of media for an additional 4.5 hours. Cells were then collected and centrifuged at 400x *g* for 5 minutes. Cell culture supernatant was carefully pipetted into a separate tube, aliquoted, and frozen at -20°C. Cells were added to a 96-well plate and stained for flow as outlined above. After surface staining, intracellular cytokines were stained using the FoxP3 Transcription Factor Buffer Set (BD Biosciences) according to the manufacturer’s instructions. The following antibodies were used: IFN*γ* (Clone: 4S.B3; Biolegend) and TNF (Clone: MAb11; Biolegend). Afterwards cells were centrifuged at 400 x *g* for 6 minutes, washed once with Fix/Perm wash, once with FACS buffer, and resuspended in 200 μL FACS buffer for analysis.

### Seahorse Assay

Glycolysis in expanded NK cells was measured with a Glycolytic Rate Assay kit (Agilent, Santa Clara, CA) following the manufacturer’s instructions using a Seahorse XFe96 Analyzer. Assays were run with replicates of 6 at a density of 4 x 10^5^ cells per well.

### Enzyme Linked Immunosorbent Assay (ELISA)

Cytokines were detected with either the Human IFN*γ* ELISA Kit Picokine or the Human TNF ELISA Kit Picokine (Booster Bio, Pleasanton, CA). ELISAs were performed according to the manufacturer’s instructions. Cell culture supernatants were diluted 1:100 for IFN*γ* ELISA and 1:10 for the TNF ELISA. Freshly thawed cell culture supernatants were used for each run of the ELISA and supernatants were not freeze-thawed multiple times before running.

### Electroporation of CRISPR RNPs into NK Cells

NK cells were collected and centrifuged at 400x *g* for 5 minutes, washed in PBS, centrifuged again, and resuspended at ~1.25 x 10^8^ cells / mL (i.e., 2.5 x 10^6^ cells in a volume of 20 μL) in Electroporation Buffer (MaxCyte, Rockville, MD). Multiplexed sgRNA’s ordered from Synthego (Redwood City, CA) were resuspended at 100 μM in sterile water. The sgRNA guides sequences were: *ZBTB16* (CAGAACCCUAGCCACCCCAC), *KLRC1* (UGAACAGGAAAUAACCUAUG), *EPAS1* (GAUUGCC-AGUCGCAUGAUGG, UUAGGAGUAGCUCGGAGAGG, GACGGAGGUGUUCUAUGAGC). The following PCR (forward: GGTGTGAGCCCAGATCAGTC; Reverse: TTCTGTGGGTGAAAGTTCTGT) and sequencing (GGTGTGAGCCCAGATCAGTCTAGTAAG) primers were used for EPAS1. Next, 3 μL of Alt-R™ S.p. HiFi Cas9 Nuclease V3 (IDT, Newark, NJ) was combined with 3.75 μL of multiplexed sgRNA’s targeting *EPAS1* and complexed for 15-30 minutes at room temperature (“RNP mixture”). Next, 20 μL of cell suspension was added to the wells of a MaxCyte OC 25×3 processing assembly (PA) before adding 5.9 μL of RNP mixture. PAs were then electroporated using either the NK-4 or NK-6 programs on a MaxCyte ATx before resting in a 37°C incubator for 30 minutes. Finally, cells were carefully removed from their PA and replated in pre-warmed complete RPMI-1640 supplemented with 600 U/mL of human IL-2. DNA was extracted after 48 hours from a portion of cells using the DNeasy Blood and Tissue kit (Qiagen) according to the manufacturer’s instructions. Following amplification using primer sequences provided by Synthego and purification of the PCR reaction, the resulting DNA was sent for Sanger sequencing at the CCHMC Genomics Sequencing Facility. KO efficiency was calculated using Synthego’s Inference of CRISPR Edits (ICE) algorithm.

### Statistical analysis

Statistical analyses were performed throughout using GraphPad Prism software (Dotmatic, Boston, MA). Each dataset was first tested for normality using the Shapiro-Wilk test. If the datasets passed the normality test, significance was assessed using a paired t test, a 2-way ANOVA, or multiple one-sample t tests with FDR correction using two-stage linear step-up procedure of Benjamini, Krieger, and Yekutieli. For nonparametric data we employed a Wilcoxon matched pairs signed rank test or a Wilcoxon signed rank test for analysis.

## RESULTS

### HIF-2α is required for ex vivo maintenance of activated human NK cells

Past studies suggest that hypoxia predominately modulates the antitumor capacity of NK cells and CD8^+^ T cells via HIF-1α-dependent mechanisms (23, 33). The role of HIF-2α (encoded by *EPAS1*) in cytotoxic lymphocytes remains unclear. However, forced overexpression of HIF-2α, but not HIF-1α, in the context of normoxia increases the cytotoxic differentiation and antitumor cytolytic function of mouse CD8 T cells (34). Corresponding CRISPR-mediated ablation of *Epas1* promoted terminal exhaustion and a reduced capacity for tumor cell killing in mouse CD8 T cells (35). The potential for HIF-2α to contribute to the antitumor capacity of NK cells, and more generally the cytolytic capacity of human immune cells, remains poorly defined.

Towards this goal, we first adapted a protocol for CRISPR-mediated editing of primary human NK cells (36). We validated our technical capabilities for gene perturbation in these cells by targeting the cell surface receptor NKG2A (encoded by *KLRC1*) and transcription factor PLZF (encoded by *ZBTB16*). Using the NK4 electroporation program on a Maxcyte ATx, we delivered a validated *ZBTB16* sgRNA (37) and mRNA encoding *Cas9* (38) to primary human NK cells. We observed dose-dependent 29% (50ng sgRNA) and 57% (100ng sgRNA) reductions in expression of PLZF on day 4 of culture after electroporation (**Figure 1A**). This was accompanied by a minimal loss (6%) of cell viability (**Figure 1B**). In a similar manner, we delivered Cas9-sgRNA ribonucleoprotein (RNP) complexes targeting *KLRC1* (39) to primary human NK cells and observed a mean 76% reduction in NKG2A expression with minimal loss of viability (**Figure 1C**). These results demonstrate our reproducible capacity to target genes in primary NK cells for CRISPR perturbation without appreciable cell death caused by the experimental procedures.

**Figure 1.**
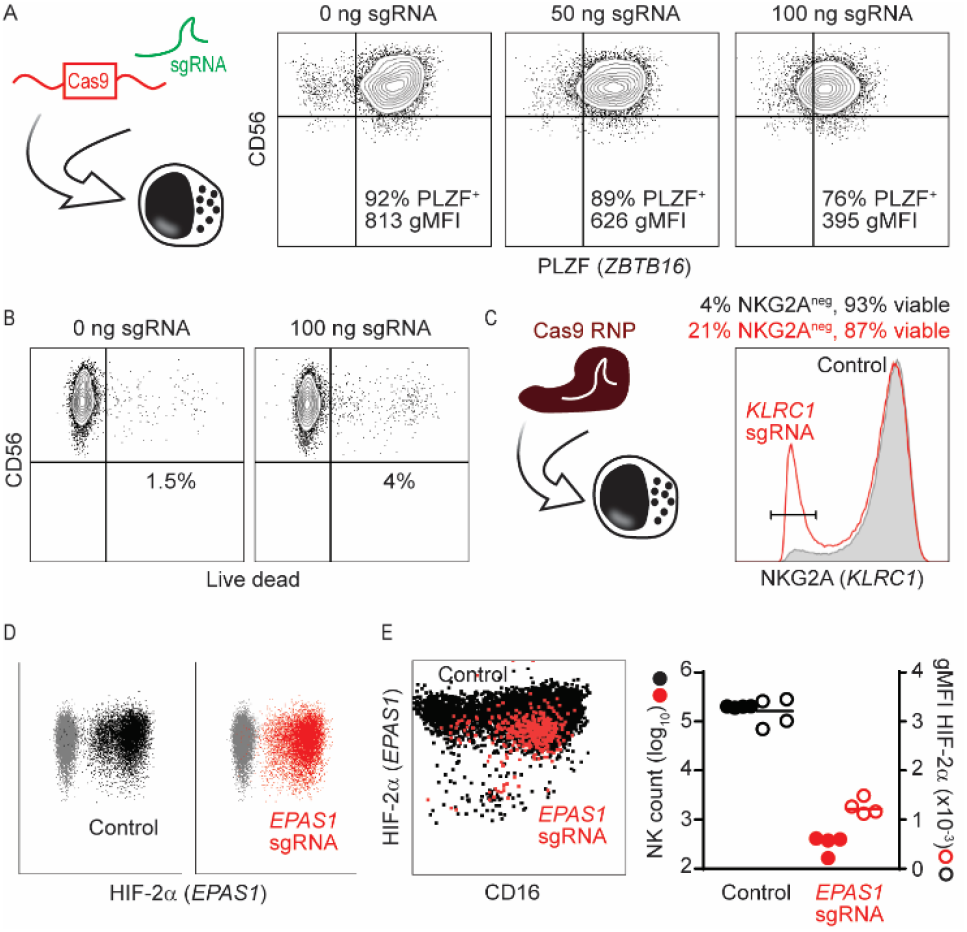
CRISPR-Cas9 editing of *EPAS1* decreases NK cell persistence. (**A**) CRISPR-Cas9 editing of primary human NK cells cultured in IL-2 via electroporation with sgRNA encoding Cas9 protein and sgRNA targeting *ZBTB16* (PLZF). Representative flow cytometry staining on day 4 post-electroporation shows (**A**) reduced proportions of PLZF+ NK cells and geometric mean fluorescence intensity (gMFI) expression levels of PLZF with increasing doses of sgRNA targeting *ZBTB16*, and (**B**) high viability of NK cells electroporated with and without sgRNA cargo. (**C**) Cas9 ribonucleoprotein (Cas9 RNP) editing in primary human NK cells to knock down *KLRC1* (NKG2A, red curve) reduces the expression of NKG2A relative to electroporation of Cas9 without sgRNA (gray curve). Mean % of viable NK cells and NKG2A-negative NK cells labeled for each group as determined by flow cytometry. (**D**) Dot plots compare primary human NK cells electroporated with Cas9 RNP targeting *EPAS1* and stained for HIF-2α expression after 6 days. (**E**) 24 hours after electroporation with GFP-Cas9 RNP targeting *EPAS1*, primary NK cells were sorted into GFP^-^ (black) or GFP^+^ (red) populations and re-analyzed after an additional 5 days of culture. Flow cytometry staining for viable NK cells within these populations is shown on the left. gMFI of HIF-2α and NK cell enumeration are shown on right.

To assess the role of HIF-2α in primary NK cells, we next electroporated NK cells with multiplexed *EPAS1*-targeting Cas9 RNP complexes. Unlike prior CRISPR experiments, we did not observe any change in HIF-2α expression after targeting *EPAS1* in the resulting pool of expanded NK cells after four days of culture through multiple experimental replicates (**Figure 1D**). Sequencing analysis using the Inference of CRISPR Edits (ICE) tool revealed an exceedingly low efficiency of editing in expanded pool of edited NK cells, despite EditCo’s (formerly Synthego) proprietary XDel algorithm, as well as CRISPOR (40) and CHOPCHOP (41), predicting comparable KO efficiency for our chosen guides compared to other published guides. To determine if *EPAS1*-edited cells might be less fit and out-competed by unedited cells in the culture, we repeated these experiments using a GFP-conjugated Cas9 to facilitate sorting of GFP^+^ (*Cas9*-electroporated) NK cells. Twenty-four hours after electroporation, GFP^+^ and GFP^-^ NK cells were sorted and cultured separately for an additional five days. Expanded GFP^-^ NK cells exhibited high viability (85%) and characteristic expression of CD16 and CD56 (**Figure 1E**). In contrast, cultures of GFP^+^ NK cells contained few cells and exhibited low viability (**Figure 1E**). Notably, decreased viability was not observed in electroporation controls (no Cas9 or RNP) or following delivery of Cas9 alone to cells (data not shown). While these outcomes complicated validation of selective editing of *EPAS1*, the results contribute to a hypothesis that HIF-2α is important for persistence or fitness of expanded human NK cells.

### Cytokine-driven accumulation of HIF-2α in NK cells during normoxia

The cytokines IL-2 and IL-15 are used to promote proliferative expansion and survival of NK cells during *ex vivo* cultures and for maintenance of therapeutic NK cells in patients (42, 43). In most cell types exposed to standard oxygen concentrations (18-20% O_2_), HIF-1α and HIF-2α protein expression is restricted via rapid hydroxylation, ubiquitination, and proteasomal degradation (17). Therefore, we hypothesized that IL-2 and IL-15 must stabilize or promote HIF-2α protein expression in primary NK cells. In addition to increased mRNA expression of *EPAS1* after cytokine treatment (**Figure 2A**), culture of primary human NK cells in IL-2 in the context of normal oxygen levels resulted in increased protein expression of HIF-2α (**Figure 2B,C**). In contrast, HIF-1α levels were not changed by *ex vivo* culture with these cytokines (**Figure 2C**), consistent with prior work demonstrating the HIF-1α stabilization in NK cells requires hypoxic conditions (28). Culture of NK cells in IL-15 (**Figure 2D**) and activation of NK cells with a combination of IL-12 and IL-18 (**Figure 2E**) also increased expression of HIF-2α protein. Collectively, these data suggest that cytokines that stimulate proliferation, survival and effector functions of NK cells also selectively induce high expression levels of HIF-2α in the absence of hypoxia.

**Figure 2.**
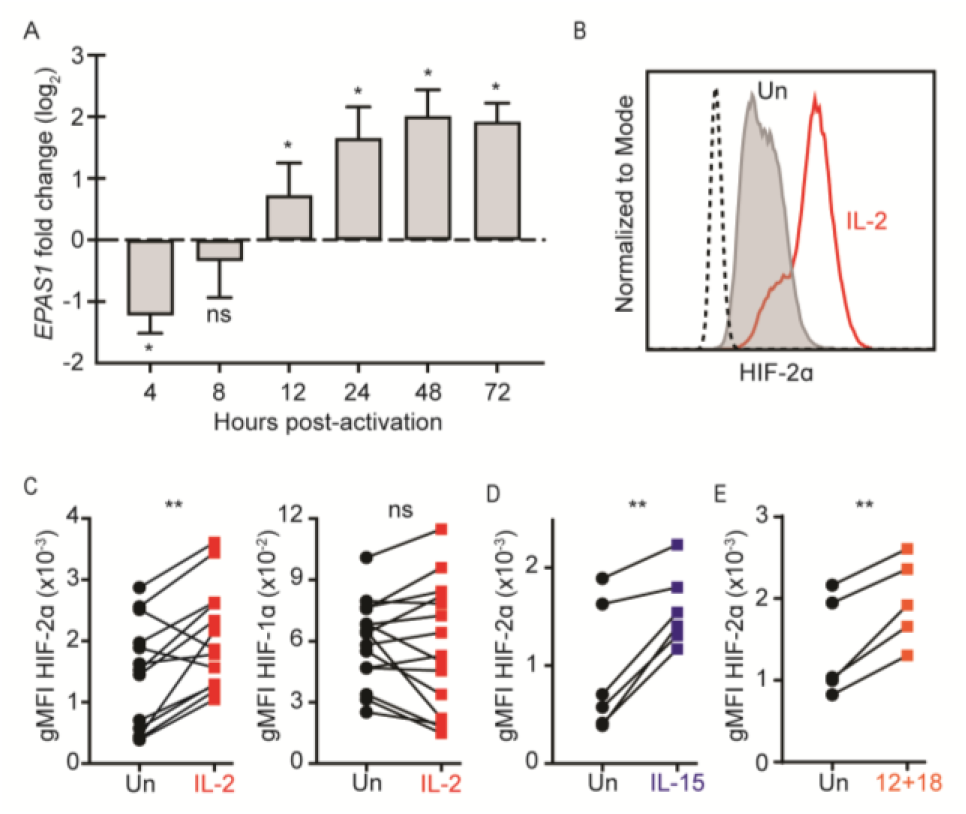
NK cell expression of HIF-2α increases following cytokine stimulation. Primary human NK cells were stimulated with (**A-C**) IL-2, (**D**) IL-15, or (**E**) IL-12 and IL-18. (**A**) Mean (± standard deviation) normalized expression of *EPAS1* RNA determined by qPCR at various times after IL-2 stimulation relative to baseline expression (n = 6 donors). Statistical significance was determined using Wilcoxon signed-rank test with FDR correction set to 1%. (**B**) Representative histogram showing HIF-2α protein expression in unstimulated (Un) NK cells (grey) or after 3 days after IL-2 treatment (red) relative to isotype control staining (dotted black). (**C**) Geometric MFI (gMFI) of HIF-2α (left) or HIF-1α (right) expression in unstimulated (Un) or IL-2-stimulated NK cells on day 3 of culture (n = 15 donors) (**D**,**E**) gMFI of HIF-2α expression in unstimulated (Un) NK cells relative to (**D**) day 3 of culture in IL-15 or (**E**) after 36 hours of culture with IL-12 and IL-18 (n = 5 donors). Statistically significant differences between groups in C-E were determined by paired t test. *ns*, p>0.05, *, p<0.05, **, p<0.01

### Classical *HIF-2α-HIF-1β transcriptional activity is dispensable in human NK cells*

To assess the consequences of HIF-2α expression in NK cells, we leveraged a well-validated inhibitor of the conventional transcriptional functions of HIF-2α, PT2385. This inhibitor blocks heterodimerization of HIF-2α and HIF-1β to prevent transcriptional activity at hypoxia responsive elements throughout the genome (44-46). PT2385 CAY-10585 altered either the baseline or maximum rate of glycolysis in expanded NK cells compared to vehicle-treated NK cells (**Figure 3D**). Collectively, our experimental data suggest that despite high expression of HIF-2α in activated NK cells, inhibition of classical HIF-2α/HIF-1β transcriptional activities does not dramatically impact NK cell effector function.

**Figure 3.**
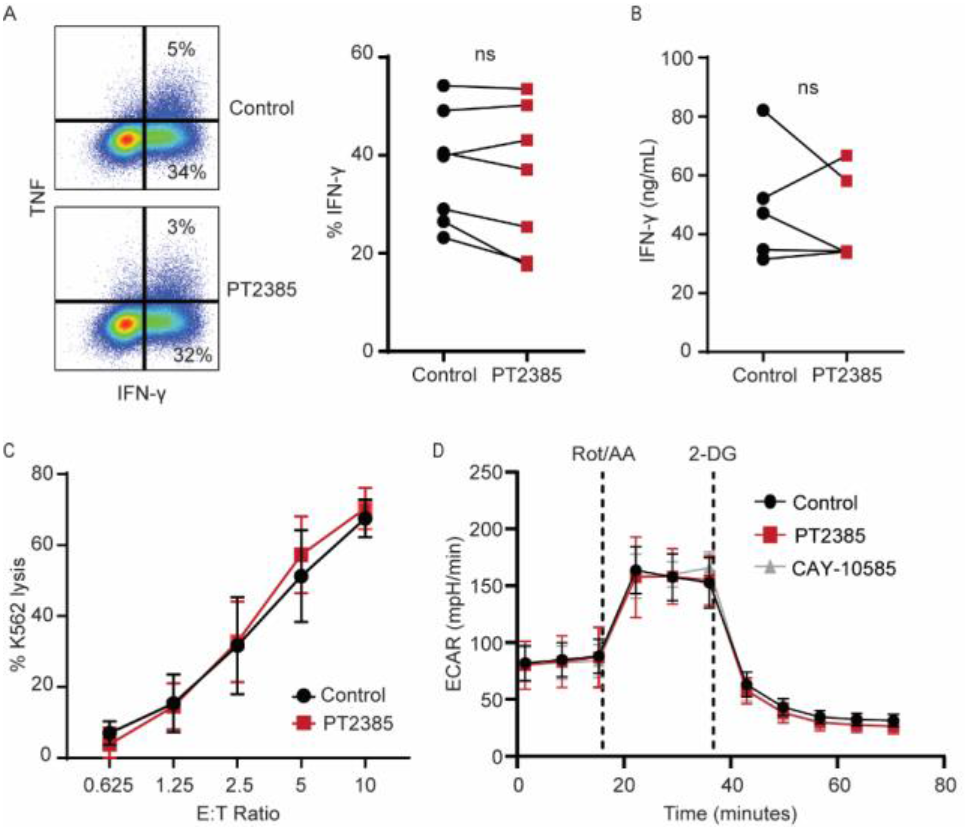
Classical HIF-2α/HIF-1β transcriptional activity does not impact NK cell function. Primary human NK cells were treated with either DMSO (control) or an inhibitor of HIF-2α/HIF-1β heterodimer transcriptional activity (PT2385) in the presence of IL-12 and IL-18 for 20 hours. Brefeldin A and monensin were added for the final 4 hours of culture. (**A**) Representative gating of IFN*γ* and TNF expressing NK cells is shown (left) with graph indicating the mean proportion of IFN-*γ*^+^ NK cells across 7 donors (right). (**B**) IFN-*γ* in supernatants of NK cells treated with IL-12/IL-18 and DMSO (control) or PT2385 (n = 5 donors). (**C**) Lysis of K562 target cells following co-culture with NK cells treated with or without PT2385 at different effector to target ratios (n = 4 donors). (**D**) NK cells were expanded using CTS Expander for 10 days after isolation. The graph shows extracellular acidification rate (ECAR) as a measure of glycolytic rate in cells treated with DMSO (control), HIF-2α/HIF-1β inhibitor (PT2385), or HIF-1α inhibitor (CAY-10585) (n = 5-6 donors). The lack of statistically significant differences between groups were assessed by a (**A**) paired t test, (**B**) Wilcoxon matched-pairs signed rank test, or (**C**,**D**) 2-way ANOVA.

### HIF-2α is required for fitness and persistence of NK cells

To address the disparity between the requirement for HIF-2α in our CRISPR editing experiments and the apparent dispensability of HIF-2α/HIF-1β activity in NK cells, we employed compound 76 (C76) as a small molecule inhibitor that blocks HIF-inhibition of HIF-2α/HIF-1β had no measurable effect on IFN-*γ* expression by NK cells stimulated with IL-12 and IL-18 (**Figure 3A**). Likewise, the quantities of IFN-*γ* secreted by IL-12/IL-18-activated NK cells were not affected by PT2385 (**Figure 3B**). In addition, there was no statistically significant change in the capacity of NK cells to kill prototypical K562 tumor target cells following treatment with PT2385 (**Figure 3C**). Thus, the functional capacity of activated human NK cells is not a measurable downstream target of conventional HIF-2α transcriptional activity.

We next considered the potential impact of HIF-2α on NK cell metabolism. Of note, activated NK cells are reliant on glycolysis (47) and HIF-2α is reported to increase the glycolytic capacity of certain tumor cells (48). However, neither HIF-2α/HIF-1β inhibition using PT2385 nor HIF-1α inhibition with 2α protein translation (49). C76 increases the binding of Iron Regulatory Protein 1 (IRP1) to the iron responsive element (IRE) in the 5’ UTR of *EPAS1* mRNA to block access of the small ribosomal subunit and thereby decrease translation (50). Treatment of cytokine-stimulated NK cells with C76 reduced the expression of HIF-2α protein by roughly two-fold on days 3 and 4 of culture (**Figure 4A**). Similar to CRISPR perturbation of *EPAS1*, C76 ablation of HIF-2α resulted in substantially reduced cellularity of the cultures and a loss of cells with features (CD56 and CD16 expression) of NK cells (**Figure 4B**). As in Figure 3, inhibition of HIF-2α/HIF-1β heterodimerization and classical transcriptional function of HIF-2α with PT2385 had no apparent effect on NK cells in these cultures (**Figure 4B**). An inhibitor of HIF-1α translation (KC7F2) (51) also did not exert an appreciable impact on NK cells (**Figure 4B**). C76 most consistently affected the persistence of CD56^dim^ CD16^+^ cells, while CD56^bright^ CD16^-^ NK cells were less directly affected (**Figure 4C**). Notably, cultures of CD3^+^ T cells and CD19^+^ B cells were minimally impacted by treatment with C76 (**Figure 4D**). Thus, the reduction of HIF-2α expression with C76 is not inherently toxic to lymphocytes but may be selectively detrimental to NK cells. Indeed, treatment with C76 increased the reactivity of NK cells with Annexin V, consistent with exposure of phosphatidylserine during cell death (**Figure 4E**). Therefore, the expression of HIF-2α in cytokine-activated human NK cells selectively promotes the survival and persistence of these lymphocytes independently of canonical HIF-2α/HIF-1β transcription factor activity.

**Figure 4.**
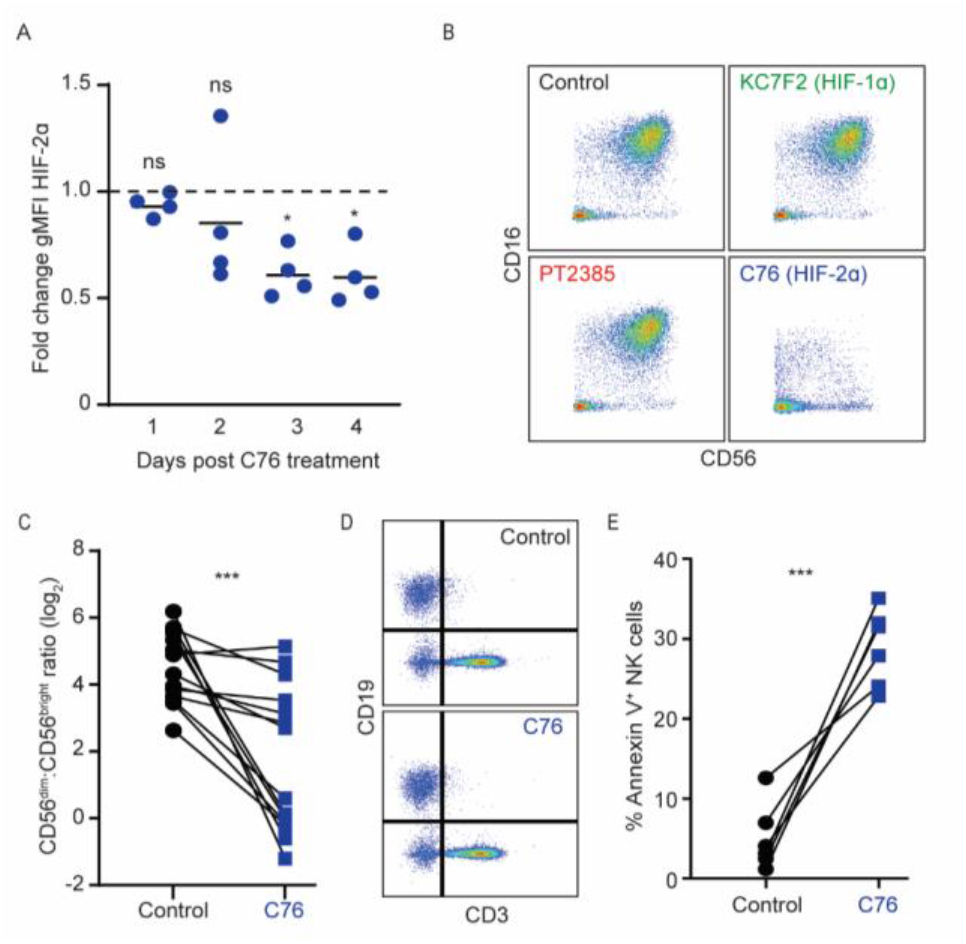
HIF-2α protein supports NK cell fitness during *in vitro* culture. (**A**) Reduced protein expression of HIF-2α on NK cells determined by flow cytometry gMFI at multiple time points after culture in IL-2 and treatment with HIF-2α translational inhibitor (Compound 76, C76) normalized to DMSO-treated time-matched controls (dashed line). Statistically significant differences were determined using multiple one-sample t tests with FDR correction set to 1% (n = 4 donors). (**B**) Representative pseudocolor plots showing remaining NK cells on day 4 of cultures treated with DMSO (Control), inhibitor of HIF-2 α/HIF-1β heterodimerization (PT2385), inhibitor of HIF-1α translation (KC7F2), or inhibitor of HIF-2α translation (C76). (**C**) Ratio CD56^dim^ NK cells to CD56^bright^ NK cells on day 4 of culture with DMSO (Control) or C76. Statistical significance was determined using a Wilcoxon matched pairs signed rank test (n = 14 donors). (**D**) Representative plots showing persistence of T (CD3) and B (CD19) cells after 4 days of treatment with C76. (**E**). Proportion of Annexin V^+^ NK cells on day 3 after C76 treatment with differences assessed using paired t test (n = 6 donors). *ns*, p>0.05, *, p<0.05, ***, p<0.001

## DISCUSSION

Our study provides the first evidence of a role for HIF-2α in NK cells. Perturbation of HIF-2α expression via CRISPR or translation inhibitors independently reveal a survival advantage conferred by HIF-2α on cytokine-activated human NK cells. This is consistent with elevated expression of HIF2α but not HIF-1α following cytokine-activation of NK cells. Importantly, the pro-survival role of HIF-2α in cytokine-activated NK cells is not shared by HIF-1α and is independent of reduced oxygen availability. Moreover, this activity is distinct from conventional regulation of hypoxia-responsive gene expression through transcriptional activities of HIF-2α/HIF-1β heterodimers. Collectively, these data establish a unique and unconventional contribution of HIF-2α to survival and persistence of NK cells during inflammation.

The stabilization and increased expression of hypoxia inducible factors in response to limited oxygen availability enables the expression of gene programs for cells to adapt to hypoxic conditions (~1% O_2_). Investigation of NK cell responses to hypoxia, particularly in the context of solid tumors, has largely focused on the role of the HIF-1α isoform with little clarity regarding the contributions, if any, of HIF-2α (23-30, 52). Although HIF-1α expression by NK cells during hypoxia can be amplified by cytokines, cytokine stimulation alone could not induce HIF-1α protein expression in the context of standard normoxia (28, 30). In contrast, we show that HIF-2α but not HIF-1α is up-regulated by NK cells in response to the cytokines IL-2, IL-15, and IL-12/IL-18. Thus, HIF-2α stabilization may represent a parallel adaptation of NK cells that facilitates gene expression programs benefiting NK cells during inflammation.

Given the vital role of Signal Transducers and Activators of Transcription (STAT) proteins in the responses of NK cells to IL-2, IL-15, and IL-12, these proteins are likely candidates underlying the transcriptional upregulation of *EPAS1* in response to these cytokines. The IL-2 and IL-15 receptors primarily signal via STAT5A and STAT5B (53, 54). Indeed, STAT5 binds directly to the *EPAS1* promoter in hematopoietic stem cells (55). Moreover, expression of activating mutants of *STAT5B* in human NK cell lines results in enrichment of STAT5 at the *EPAS1* promoter (56). Although IL-12 primarily activates STAT4, which is not appreciably enriched at the *EPAS1* promoter of IL-12-stimulated mouse NK cells (57), IL-12 can concomitantly stimulate STAT5 activation (58, 59). Alternatively, HIF-2α has also been reported as a downstream target of PI3K-mTOR signaling complex (60, 61), which is also a component of the signaling cascade downstream of common gamma chain cytokines (42). Future studies are needed to reveal the precise transcriptional networks involved in increased transcription of *EPAS1* in cytokine-activated NK cells. Critically, the mechanisms by which cytokine stimulation permits stabilization of HIF-2α protein in the context of standard normoxia (~20% O_2_) remain ill-defined.

Consistent with the normoxia setting in which we observe that cytokines induce HIF-2α expression by NK cells, we find that classical functions of HIF-2α/HIF-1β heterodimers in enabling gene expression adaptations to hypoxia are not required for the benefits conferred by HIF-2α upon cytokine-activated NK cells. Moreover, the ability of HIF-2α to heterodimerize with HIF-1β was dispensable for a variety of NK cell functional measures the context of standard normoxia (~20% O_2_), including cytokine production and tumor cell killing. This is consistent with other reports suggesting that the conventional functions of HIF-2α/HIF-1β heterodimers are dispensable for functional responsiveness of NK cells (27, 62). While HIF-2α can regulate the expression cytotoxic mediators (*Prf1, Gzmb*), cytokine receptors (*Il2ra*), and costimulatory molecules (*Tnfrsf8*) in mouse CD8 T cells (34, 63), the function of HIF-2α may differ in NK cells due to distinct experimental contexts, cell type specific gene regulatory circuits (64), disparate expression of relevant transcription factors (65), differential chromatin accessibility (66), or dissimilar cis-regulatory elements between individual cell types (66, 67). For example, HIF-2α has been shown to regulate the glycolytic genes *SLC2A*3 and *ALDOA* in primary acute myeloid leukemia progenitor cell samples but does not regulate the same genes in CD34-expressing progenitor cells in cord blood (48). Alternative pathways of HIF-2α-mediated regulation of gene expression via heterodimerization with BMAL1 in heart tissue (68) or AP2α in melanoma cells (69), highlights the possibility of unique transcriptional partnerships by which HIF-2α governs NK cell survival following cytokine stimulation.

Among lymphocytes, the loss of HIF-2α appears to be uniquely detrimental in NK cells, whereas HIF-2α was dispensable for T and B cell survival. In fact, CRISPR-Cas9 deletion of *EPAS1* in primary T cells did not lead to substantial transcriptional changes (70). Nevertheless, HIF-2α maintains a protective role in other cell types by suppressing apoptosis (71, 72) and ferroptosis (73). To this effect, NK cells treated with an inhibitor of HIF-2α translation showed increased cell surface exposure of phosphatidylserine. Preliminary studies with an array of inhibitors of apoptosis and ferroptosis failed to rescue NK cells in the absence of HIF-2α, confounding efforts to determine the manner of cell death prevented by HIF-2α induction in cytokine activated NK cells.

In summary, our data reveal a cytokine-mediated, oxygen-independent induction of HIF-2α but not HIF-1α, expression that preserves the fitness and persistence of cytokine activated human NK cells. We demonstrate that this function is unique to NK cells among several types of human lymphocytes and is distinct from conventional HIF-2α mediated gene regulatory mechanisms. Targeting of this previously unknown activity of HIF-2α may represent a path to improve cellular therapies involving cytokine-activated NK cells by addressing clinical needs for durable persistence of NK cells. Moreover, the impact of HIF-2α expression on NK cell biology during inflammation may be relevant to human diseases where HIF-2α is known to play a pathogenic role (74).

## Acknowledgements

We thank Laura Canaday, Eshana Mukhopadhyay, Hilal Cevik, and Nora Lakes for their assistance in completing experiments. We are grateful to George Deepe and Maria Czyzyk-Krzeska for advice and reagents concerning hypoxia inducible factors. We are grateful to Senad Divanovic and Grant Schulert for thoughtful suggestions regarding experimental design. We thank MaxCyte technical specialists, especially Nitin Kularkani for providing experimental guidance. We thank Danielle Dauphers for editing the manuscript. All flow cytometric data were acquired using equipment maintained by the Research Flow Cytometry Facility in the Division of Rheumatology at Cincinnati Children’s Hospital Medical Center. The graphical abstract was created using BioRender software.

## Author contributions

Conceptualization and design of study (HAF, SDR, SNW), project supervision (SNW), acquisition of funding (LCK, MTW, HIB, SNW), performance of experiments and data acquisition (HAF, KA, KLE, SDR), data analysis (HAF, AV, SNW, KMK), drafting of the manuscript (HAF, SNW), and critical editing of the manuscript (AV, KA, KLE, SDR, HIB, LCK, MTW, KMK).

## REFERENCES

1. Bjorkstrom, N. K., B. Strunz, and H. G. Ljunggren. 2022. Natural killer cells in antiviral immunity. Nat Rev Immunol 22: 112–123.

2. Huntington, N. D., J. Cursons, and J. Rautela. 2020. The cancer-natural killer cell immunity cycle. Nat Rev Cancer 20: 437–454.

3. Anukanth, A., and S. N. Waggoner. 2026. Unconventional functions of natural killer cells in homeostasis and disease. J Immunol 215.

4. Mujal, A. M., R. B. Delconte, and J. C. Sun. 2021. Natural Killer Cells: From Innate to Adaptive Features. Annu Rev Immunol 39: 417–447.

5. Maskalenko, N. A., D. Zhigarev, and K. S. Campbell. 2022. Harnessing natural killer cells for cancer immunotherapy: dispatching the first responders. Nat Rev Drug Discov.

6. Chiossone, L., P. Y. Dumas, M. Vienne, and E. Vivier. 2018. Natural killer cells and other innate lymphoid cells in cancer. Nat Rev Immunol 18: 671–688.

7. Miller, J. S. 2013. Therapeutic applications: natural killer cells in the clinic. Hematology Am Soc Hematol Educ Program 2013: 247–253.

8. Neelapu, S. S., S. Tummala, P. Kebriaei, W. Wierda, C. Gutierrez, F. L. Locke, K. V. Komanduri, Y. Lin, N. Jain, N. Daver, J. Westin, A. M. Gulbis, M. E. Loghin, J. F. de Groot, S. Adkins, S. E. Davis, K. Rezvani, P. Hwu, and E. J. Shpall. 2018. Chimeric antigen receptor T-cell therapy - assessment and management of toxicities. Nat Rev Clin Oncol 15: 47–62.

9. Lu, H., X. Zhao, Z. Li, Y. Hu, and H. Wang. 2021. From CAR-T Cells to CAR-NK Cells: A Developing Immunotherapy Method for Hematological Malignancies. Front Oncol 11: 720501.

10. Klingemann, H. 2015. Challenges of cancer therapy with natural killer cells. Cytotherapy 17: 245–249.

11. Dagher, O. K., and A. D. Posey, Jr. 2023. Forks in the road for CAR T and CAR NK cell cancer therapies. Nature immunology 24: 1994–2007.

12. Andrea, A. E., A. Chiron, G. Sarrabayrouse, S. Bessoles, and S. Hacein-Bey-Abina. 2024. A structural, genetic and clinical comparison of CAR-T cells and CAR-NK cells: companions or competitors? Frontiers in immunology 15: 1459818.

13. Fehniger, T. A., M. A. Cooper, and M. A. Caligiuri. 2002. Interleukin-2 and interleukin-15: immunotherapy for cancer. Cytokine Growth Factor Rev 13: 169–183.

14. Denman, C. J., V. V. Senyukov, S. S. Somanchi, P. V. Phatarpekar, L. M. Kopp, J. L. Johnson, H. Singh, L. Hurton, S. N. Maiti, M. H. Huls, R. E. Champlin, L. J. Cooper, and D. A. Lee. 2012. Membrane-bound IL-21 promotes sustained ex vivo proliferation of human natural killer cells. PLoS One 7: e30264.

15. Lapteva, N., A. G. Durett, J. Sun, L. A. Rollins, L. L. Huye, J. Fang, V. Dandekar, Z. Mei, K. Jackson, J. Vera, J. Ando, M. C. Ngo, E. Coustan-Smith, D. Campana, S. Szmania, T. Garg, A. Moreno-Bost, F. Vanrhee, A. P. Gee, and C. M. Rooney. 2012. Large-scale ex vivo expansion and characterization of natural killer cells for clinical applications. Cytotherapy 14: 1131–1143.

16. LaGory, E. L., and A. J. Giaccia. 2016. The ever-expanding role of HIF in tumour and stromal biology. Nat Cell Biol 18: 356–365.

17. Semenza, G. L. 2014. Oxygen sensing, hypoxia-inducible factors, and disease pathophysiology. Annu Rev Pathol 9: 47–71.

18. Kaelin, W. G. 2005. Proline hydroxylation and gene expression. Annu Rev Biochem 74: 115–128.

19. Smythies, J. A., M. Sun, N. Masson, R. Salama, P. D. Simpson, E. Murray, V. Neumann, M. E. Cockman, H. Choudhry, P. J. Ratcliffe, and D. R. Mole. 2019. Inherent DNA-binding specificities of the HIF-1alpha and HIF-2alpha transcription factors in chromatin. EMBO Rep 20.

20. Bakleh, M. Z., and A. Al Haj Zen. 2025. The Distinct Role of HIF-1alpha and HIF-2alpha in Hypoxia and Angiogenesis. Cells 14.

21. Shi, Y., and D. M. Gilkes. 2025. HIF-1 and HIF-2 in cancer: structure, regulation, and therapeutic prospects. Cellular and molecular life sciences : CMLS 82: 44.

22. Messai, Y., M. Z. Noman, B. Janji, M. Hasmim, B. Escudier, and S. Chouaib. 2015. The autophagy sensor ITPR1 protects renal carcinoma cells from NK-mediated killing. Autophagy: 0.

23. Ni, J., X. Wang, A. Stojanovic, Q. Zhang, M. Wincher, L. Buhler, A. Arnold, M. P. Correia, M. Winkler, P. S. Koch, V. Sexl, T. Hofer, and A. Cerwenka. 2020. Single-Cell RNA Sequencing of Tumor-Infiltrating NK Cells Reveals that Inhibition of Transcription Factor HIF-1alpha Unleashes NK Cell Activity. Immunity 52: 1075–1087 e1078.

24. Krzywinska, E., C. Kantari-Mimoun, Y. Kerdiles, M. Sobecki, T. Isagawa, D. Gotthardt, M. Castells, J. Haubold, C. Millien, T. Viel, B. Tavitian, N. Takeda, J. Fandrey, E. Vivier, V. Sexl, and C. Stockmann. 2017. Loss of HIF-1alpha in natural killer cells inhibits tumour growth by stimulating non-productive angiogenesis. Nat Commun 8: 1597.

25. Victorino, F., T. M. Bigley, E. Park, C. H. Yao, J. Benoit, L. P. Yang, S. J. Piersma, E. J. Lauron, R. M. Davidson, G. J. Patti, and W. M. Yokoyama. 2021. HIF1alpha is required for NK cell metabolic adaptation during virus infection. eLife 10.

26. Nakazawa, T., T. Morimoto, R. Maeoka, K. Yamada, R. Matsuda, M. Nakamura, F. Nishimura, S. Yamada, Y. S. Park, T. Tsujimura, and I. Nakagawa. 2024. Characterization of HIF-1alpha Knockout Primary Human Natural Killer Cells Including Populations in Allogeneic Glioblastoma. Int J Mol Sci 25.

27. Pelletier, A., E. Nelius, Z. Fan, E. Khatchatourova Alvarado-Diaz, J. He, E. Krzywinska, M. Sobecki, S. Nagarajan, Y. Kerdiles, J. Fandrey, D. Gotthardt, V. Sexl, K. de Bock, and C. Stockmann. 2023. Resting natural killer cell homeostasis relies on tryptophan/NAD(+) metabolism and HIF-1alpha. EMBO Rep 24: e56156.

28. Cluff, E., C. C. Magdaleno, E. Fernandez, T. House, S. Swaminathan, A. Varadaraj, and N. Rajasekaran. 2022. Hypoxia-inducible factor-1 alpha expression is induced by IL-2 via the PI3K/mTOR pathway in hypoxic NK cells and supports effector functions in NKL cells and ex vivo expanded NK cells. Cancer Immunol Immunother 71: 1989–2005.

29. Lim, S. A., Y. Moon, M. H. Shin, T. J. Kim, S. Chae, C. Yee, D. Hwang, H. Park, and K. M. Lee. 2021. Hypoxia-Driven HIF-1alpha Activation Reprograms Pre-Activated NK Cells towards Highly Potent Effector Phenotypes via ERK/STAT3 Pathways. Cancers (Basel) 13.

30. Coulibaly, A., S. Y. Velasquez, N. Kassner, J. Schulte, M. V. Barbarossa, and H. A. Lindner. 2021. STAT3 governs the HIF-1alpha response in IL-15 primed human NK cells. Sci Rep 11: 7023.

31. Sobecki, M., E. Krzywinska, S. Nagarajan, A. Audige, K. Huynh, J. Zacharjasz, J. Debbache, Y. Kerdiles, D. Gotthardt, N. Takeda, J. Fandrey, L. Sommer, V. Sexl, and C. Stockmann. 2021. NK cells in hypoxic skin mediate a trade-off between wound healing and antibacterial defence. Nature communications 12: 4700.

32. Velasquez, S. Y., D. Killian, J. Schulte, C. Sticht, M. Thiel, and H. A. Lindner. 2016. Short Term Hypoxia Synergizes with Interleukin 15 Priming in Driving Glycolytic Gene Transcription and Supports Human Natural Killer Cell Activities. The Journal of biological chemistry 291: 12960–12977.

33. McGettrick, A. F., and L. A. J. O’Neill. 2020. The Role of HIF in Immunity and Inflammation. Cell Metab 32: 524–536.

34. Velica, P., P. P. Cunha, N. Vojnovic, I. P. Foskolou, D. Bargiela, M. Gojkovic, H. Rundqvist, and R. S. Johnson. 2021. Modified Hypoxia-Inducible Factor Expression in CD8(+) T Cells Increases Antitumor Efficacy. Cancer Immunol Res 9: 401–414.

35. Kadyrzhanova, G., M. Tamai, S. Sarkar, R. S. Kalra, and H. Ishikawa. 2025. Aging impairs CD8 T cell responses in adoptive T-cell therapy against solid tumors. Front Immunol 16: 1484303.

36. Clara, J. A., E. R. Levy, R. Reger, S. Barisic, L. Chen, E. Cherkasova, M. Chakraborty, D. S. J. Allan, and R. Childs. 2022. High-affinity CD16 integration into a CRISPR/Cas9-edited CD38 locus augments CD38-directed antitumor activity of primary human natural killer cells. J Immunother Cancer 10.

37. Liu, W., J. M. Scott, E. Langguth, H. Chang, P. H. Park, and S. Kim. 2020. FcRgamma Gene Editing Reprograms Conventional NK Cells to Display Key Features of Adaptive Human NK Cells. iScience 23: 101709.

38. Pomeroy, E. J., J. T. Hunzeker, M. G. Kluesner, W. S. Lahr, B. A. Smeester, M. R. Crosby, C. L. Lonetree, K. Yamamoto, L. Bendzick, J. S. Miller, M. A. Geller, B. Walcheck, M. Felices, B. R. Webber, T. K. Starr, and B. S. Moriarity. 2020. A Genetically Engineered Primary Human Natural Killer Cell Platform for Cancer Immunotherapy. Mol Ther 28: 52–63.

39. Bexte, T., J. Alzubi, L. M. Reindl, P. Wendel, R. Schubert, E. Salzmann-Manrique, I. von Metzler, T. Cathomen, and E. Ullrich. 2022. CRISPR-Cas9 based gene editing of the immune checkpoint NKG2A enhances NK cell mediated cytotoxicity against multiple myeloma. Oncoimmunology 11: 2081415.

40. Concordet, J. P., and M. Haeussler. 2018. CRISPOR: intuitive guide selection for CRISPR/Cas9 genome editing experiments and screens. Nucleic Acids Res 46: W242–W245.

41. Labun, K., T. G. Montague, M. Krause, Y. N. Torres Cleuren, H. Tjeldnes, and E. Valen. 2019. CHOPCHOP v3: expanding the CRISPR web toolbox beyond genome editing. Nucleic Acids Res 47: W171–W174.

42. Mao, Y., V. van Hoef, X. Zhang, E. Wennerberg, J. Lorent, K. Witt, L. Masvidal, S. Liang, S. Murray, O. Larsson, R. Kiessling, and A. Lundqvist. 2016. IL-15 activates mTOR and primes stress-activated gene expression leading to prolonged antitumor capacity of NK cells. Blood 128: 1475–1489.

43. Sharma, A., C. N. Toepfer, T. Ward, L. Wasson, R. Agarwal, D. A. Conner, J. H. Hu, and C. E. Seidman. 2018. CRISPR/Cas9-Mediated Fluorescent Tagging of Endogenous Proteins in Human Pluripotent Stem Cells. Curr Protoc Hum Genet 96: 21 11 21-21 11 20.

44. Wehn, P. M., J. P. Rizzi, D. D. Dixon, J. A. Grina, S. T. Schlachter, B. Wang, R. Xu, H. Yang, X. Du, G. Han, K. Wang, Z. Cao, T. Cheng, R. M. Czerwinski, B. S. Goggin, H. Huang, M. M. Halfmann, M. A. Maddie, E. L. Morton, S. R. Olive, H. Tan, S. Xie, T. Wong, J. A. Josey, and E. M. Wallace. 2018. Design and Activity of Specific Hypoxia-Inducible Factor-2alpha (HIF-2alpha) Inhibitors for the Treatment of Clear Cell Renal Cell Carcinoma: Discovery of Clinical Candidate ( S)-3-((2,2-Difluoro-1-hydroxy-7-(methylsulfonyl)-2,3-dihydro-1 H-inden-4-yl)oxy)-5-fluorobenzonitrile (PT2385). J Med Chem 61: 9691–9721.

45. Courtney, K. D., J. R. Infante, E. T. Lam, R. A. Figlin, B. I. Rini, J. Brugarolas, N. J. Zojwalla, A. M. Lowe, K. Wang, E. M. Wallace, J. A. Josey, and T. K. Choueiri. 2018. Phase I Dose-Escalation Trial of PT2385, a First-in-Class Hypoxia-Inducible Factor-2alpha Antagonist in Patients With Previously Treated Advanced Clear Cell Renal Cell Carcinoma. J Clin Oncol 36: 867–874.

46. Choueiri, T. K., T. M. Bauer, K. P. Papadopoulos, E. R. Plimack, J. R. Merchan, D. F. McDermott, M. D. Michaelson, L. J. Appleman, S. Thamake, R. F. Perini, N. J. Zojwalla, and E. Jonasch. 2021. Inhibition of hypoxia-inducible factor-2alpha in renal cell carcinoma with belzutifan: a phase 1 trial and biomarker analysis. Nat Med 27: 802–805.

47. Donnelly, R. P., R. M. Loftus, S. E. Keating, K. T. Liou, C. A. Biron, C. M. Gardiner, and D. K. Finlay. 2014. mTORC1-dependent metabolic reprogramming is a prerequisite for NK cell effector function. J Immunol 193: 4477–4484.

48. Wierenga, A. T. J., A. Cunningham, A. Erdem, N. V. Lopera, A. Z. Brouwers-Vos, M. Pruis, A. B. Mulder, U. L. Gunther, J. H. A. Martens, E. Vellenga, and J. J. Schuringa. 2019. HIF1/2-exerted control over glycolytic gene expression is not functionally relevant for glycolysis in human leukemic stem/progenitor cells. Cancer Metab 7: 11.

49. Zimmer, M., B. L. Ebert, C. Neil, K. Brenner, I. Papaioannou, A. Melas, N. Tolliday, J. Lamb, K. Pantopoulos, T. Golub, and O. Iliopoulos. 2008. Small-molecule inhibitors of HIF-2a translation link its 5’UTR iron-responsive element to oxygen sensing. Mol Cell 32: 838–848.

50. Muckenthaler, M., N. K. Gray, and M. W. Hentze. 1998. IRP-1 binding to ferritin mRNA prevents the recruitment of the small ribosomal subunit by the cap-binding complex eIF4F. Mol Cell 2: 383–388.

51. Narita, T., S. Yin, C. F. Gelin, C. S. Moreno, M. Yepes, K. C. Nicolaou, and E. G. Van Meir. 2009. Identification of a novel small molecule HIF-1alpha translation inhibitor. Clin Cancer Res 15: 6128–6136.

52. Yu, X., J. Zhou, J. Lei, H. Ge, Z. Zhang, Y. Fu, X. Han, Q. Hu, H. Ding, W. Geng, H. Shang, and Y. Jiang. 2025. Impaired natural killer cell migration in HIV-infected individuals is caused by TIGIT-mediated inhibition of HIF-1alpha-dependent glycolysis. Cell Death Dis 16: 805.

53. Hou, J., U. Schindler, W. J. Henzel, S. C. Wong, and S. L. McKnight. 1995. Identification and purification of human Stat proteins activated in response to interleukin-2. Immunity 2: 321–329.

54. Lin, J. X., T. S. Migone, M. Tsang, M. Friedmann, J. Weatherbee, L. Zhou, A. Yamauchi, E. T. Bloom, J. Mietz, S. John, and et al. 1995. The role of shared receptor motifs and common Stat proteins in the generation of cytokine pleiotropy and redundancy by IL-2, IL-4, IL-7, IL-13, and IL-15. Immunity 2: 331–339.

55. Fatrai, S., A. T. Wierenga, S. M. Daenen, E. Vellenga, and J. J. Schuringa. 2011. Identification of HIF2alpha as an important STAT5 target gene in human hematopoietic stem cells. Blood 117: 3320–3330.

56. Kucuk, C., B. Jiang, X. Hu, W. Zhang, J. K. Chan, W. Xiao, N. Lack, C. Alkan, J. C. Williams, K. N. Avery, P. Kavak, A. Scuto, E. Sen, P. Gaulard, L. Staudt, J. Iqbal, W. Zhang, A. Cornish, Q. Gong, Q. Yang, H. Sun, F. d’Amore, S. Leppa, W. Liu, K. Fu, L. de Leval, T. McKeithan, and W. C. Chan. 2015. Activating mutations of STAT5B and STAT3 in lymphomas derived from gammadelta-T or NK cells. Nat Commun 6: 6025.

57. Wiedemann, G. M., E. K. Santosa, S. Grassmann, S. Sheppard, J. B. Le Luduec, N. M. Adams, C. Dang, K. C. Hsu, J. C. Sun, and C. M. Lau. 2021. Deconvoluting global cytokine signaling networks in natural killer cells. Nat Immunol 22: 627–638.

58. Wiedemann, G. M., S. Grassmann, C. M. Lau, M. Rapp, A. V. Villarino, C. Friedrich, G. Gasteiger, J. J. O’Shea, and J. C. Sun. 2020. Divergent Role for STAT5 in the Adaptive Responses of Natural Killer Cells. Cell Rep 33: 108498.

59. Shemesh, A., H. Pickering, K. T. Roybal, and L. L. Lanier. 2022. Differential IL-12 signaling induces human natural killer cell activating receptor-mediated ligand-specific expansion. J Exp Med 219.

60. Mohlin, S., A. Hamidian, K. von Stedingk, E. Bridges, C. Wigerup, D. Bexell, and S. Pahlman. 2015. PI3K-mTORC2 but not PI3K-mTORC1 regulates transcription of HIF2A/EPAS1 and vascularization in neuroblastoma. Cancer Res 75: 4617–4628.

61. Chen, J., J. Chen, J. Huang, Z. Li, Y. Gong, B. Zou, X. Liu, L. Ding, P. Li, Z. Zhu, B. Zhang, H. Guo, C. Cai, and J. Li. 2019. HIF-2alpha upregulation mediated by hypoxia promotes NAFLD-HCC progression by activating lipid synthesis via the PI3K-AKT-mTOR pathway. Aging (Albany NY) 11: 10839–10860.

62. De Waele, J., T. Verhezen, H. W. Lau, J. Melis, L. Brants, J. Van Audenaerde, J. de Beukelaar, C. Gevaerts, F. Lardon, A. Wouters, and E. Smits. 2025. HIF isoforms contribute distinctly during human NK cell activation. bioRxiv: 2025.2006.2027.661904.

63. Doedens, A. L., A. T. Phan, M. H. Stradner, J. K. Fujimoto, J. V. Nguyen, E. Yang, R. S. Johnson, and A. W. Goldrath. 2013. Hypoxia-inducible factors enhance the effector responses of CD8(+) T cells to persistent antigen. Nat Immunol 14: 1173–1182.

64. Li, J. H., A. Zhou, C. D. Lee, S. N. Shah, J. H. Ji, V. Senthilkumar, E. T. Padilla, A. B. Ball, Q. Feng, C. G. Bustillos, L. Riggan, A. Greige, A. S. Divakaruni, F. Annese, J. A. Cooley Coleman, S. A. Skinner, C. W. Cowan, and T. E. O’Sullivan. 2024. MEF2C regulates NK cell effector functions through control of lipid metabolism. Nat Immunol 25: 778–789.

65. Yang, P., R. Corbett, L. Daharsh, J. H. Uribe, K. A. Byrne, C. L. Loving, and C. Tuggle. 2024. Definition of regulatory elements and transcription factors controlling porcine immune cell gene expression at single cell resolution using single nucleus ATAC-seq. Genomics 116: 110944.

66. Fang, D., K. Cui, Y. Cao, M. Zheng, T. Kawabe, G. Hu, J. S. Khillan, D. Li, C. Zhong, D. Jankovic, A. Sher, K. Zhao, and J. Zhu. 2022. Differential regulation of transcription factor T-bet induction during NK cell development and T helper-1 cell differentiation. Immunity 55: 639–655 e637.

67. Awdeh, A., M. Turcotte, and T. J. Perkins. 2024. Identifying transcription factors with cell-type specific DNA binding signatures. BMC Genomics 25: 957.

68. Ruan, W., T. Li, I. H. Bang, J. Lee, W. Deng, X. Ma, C. Luo, F. Du, S. H. Yoo, B. Kim, J. Li, X. Yuan, K. Figarella, Y. A. An, Y. Y. Wang, Y. Liang, M. DeBerge, D. Zhang, Z. Zhou, Y. Wang, J. M. Gorham, J. G. Seidman, C. E. Seidman, S. F. Aranki, R. Nair, L. Li, J. Narula, Z. Zhao, A. A. Gorfe, J. D. Muehlschlegel, K. L. Tsai, and H. K. Eltzschig. 2025. BMAL1-HIF2A heterodimer modulates circadian variations of myocardial injury. Nature 641: 1017–1028.

69. Steunou, A. L., M. Ducoux-Petit, I. Lazar, B. Monsarrat, M. Erard, C. Muller, E. Clottes, O. Burlet-Schiltz, and L. Nieto. 2013. Identification of the hypoxia-inducible factor 2alpha nuclear interactome in melanoma cells reveals master proteins involved in melanoma development. Mol Cell Proteomics 12: 736–748.

70. Madsen, K. N., S. Kashima, H. Soulati, L. Wirth, K. Ito, Z. A. Yochum, V. Moritz, J. Walker, M. Machaalani, F. Ghali, P. Kenney, A. Adeniran, M. Hurwitz, T. K. Choueiri, and D. A. Braun. 2026. Pharmacologic HIF-2alpha inhibition preserves T cell effector function in clear cell renal cell carcinoma. J Clin Invest.

71. Barra, J., C. R. Liwski, P. Phonchareon, L. Portales-Cervantes, D. Gaston, T. K. Karakach, I. D. Haidl, and J. S. Marshall. 2025. IL-5 enhances human mast cell survival and interferon responses to viral infection. J Allergy Clin Immunol 155: 1968–1980.

72. Rouault-Pierre, K., L. Lopez-Onieva, K. Foster, F. Anjos-Afonso, I. Lamrissi-Garcia, M. Serrano-Sanchez, R. Mitter, Z. Ivanovic, H. de Verneuil, J. Gribben, D. Taussig, H. R. Rezvani, F. Mazurier, and D. Bonnet. 2013. HIF-2alpha protects human hematopoietic stem/progenitors and acute myeloid leukemic cells from apoptosis induced by endoplasmic reticulum stress. Cell Stem Cell 13: 549–563.

73. Green, Y. S., M. C. Ferreira Dos Santos, D. G. Fuja, E. C. Reichert, A. R. Campos, S. J. Cowman, K. Acuna Pilarte, J. Kohan, S. R. Tripp, E. A. Leibold, D. Sirohi, N. Agarwal, X. Liu, and M. Y. Koh. 2022. ISCA2 inhibition decreases HIF and induces ferroptosis in clear cell renal carcinoma. Oncogene 41: 4709–4723.

74. Ormiston, M. L., C. Chang, L. L. Long, E. Soon, D. Jones, R. Machado, C. Treacy, M. R. Toshner, K. Campbell, A. Riding, M. Southwood, J. Pepke-Zaba, A. Exley, R. C. Trembath, F. Colucci, M. Wills, J. Trowsdale, and N. W. Morrell. 2012. Impaired natural killer cell phenotype and function in idiopathic and heritable pulmonary arterial hypertension. Circulation 126: 1099–1109.

